# A large-scale brain network of species-specific dynamic human body perception

**DOI:** 10.1101/2022.07.22.501117

**Authors:** Baichen Li, Marta Poyo Solanas, Giuseppe Marrazzo, Rajani Raman, Nick Taubert, Martin Giese, Rufin Vogels, Beatrice de Gelder

## Abstract

This ultrahigh field 7T fMRI study addressed the question of whether there exists a core network of brain areas at the service of different aspects of body perception. Participants viewed naturalistic videos of monkey and human faces, bodies, and objects along with mosaic-scrambled videos for control of low-level features. ICA-based network analysis was conducted to find body and species modulations at both the voxel and the network levels. Among the body areas, the highest species selectivity was found in the middle frontal gyrus and amygdala. Two large-scale networks were highly selective to bodies, dominated by the lateral occipital cortex and right superior temporal sulcus (STS) respectively. The right STS network showed high species selectivity, and its significant human body-induced node connectivity was focused around the extrastriate body area (EBA), STS, temporoparietal junction (TPJ), premotor cortex, and inferior frontal gyrus (IFG). The human body-specific network discovered here may serve as a brain-wide internal model of the human body serving as an entry point for a variety of processes relying on body descriptions as part of their more specific categorization, action, or expression recognition functions.

## INTRODUCTION

Social species make extensive use of collaborative and competitive signals from conspecifics, allowing them to navigate successfully in the natural and social world. In the visual domain, social signals from faces and bodies are the central sources of information about conspecific presence, intentions, emotions, and actions. An extensive literature on face perception has already illustrated already the importance of face perception for regulating interactions between nearby conspecifics (Panksepp, 1989). Like the face, the body is a rich and powerful means of social communication allowing quick and easy inferences about identity, gender, sex, physical health, attractiveness, emotional state, and social status. Body perception operates at a much longer distance than face perception and provides information about emotions, intentions, and actions relevant for social interaction (de Gelder et al., 2010). Yet, aside from studies of the body as a perceptual object category, our understanding of whole-body perception is still very limited (de Gelder and Poyo Solanas, 2021; Taubert et al., 2022). Despite a vast literature on the perception of action and intention that in fact assumes that body perception is involved (Orban et al., 2021), recent theories about social perception and social brain networks do not yet integrate findings from body perception studies (Patel et al., 2019; Pitcher and Ungerleider, 2021). Doing so may enrich and diversify those models.

In view of the relevance of bodily communication, one may expect that preferential processing routes exist in the brain for bodies (Downing and Kanwisher, 2001) and body expressions (de Gelder et al., 2010), just as has long been assumed for faces (Gross et al., 1969). Previous studies on body perception mainly addressed body category-specific processes in the ventral stream. In human studies, body selective areas were reported in the middle occipital/temporal gyrus termed the extrastriate body area (EBA) (Downing and Kanwisher, 2001), in the fusiform cortex termed the fusiform body area (FBA) (Peelen and Downing, 2005; Schwarzlose et al., 2005) and in the posterior superior temporal sulcus (pSTS) (Candidi et al., 2015). Body patches observed in monkeys with fMRI were mainly found along the STS (Vogels, 2022). Similar to the situation in human studies, there is a consensus that these different areas or patches presumably have different computational functions, but there is currently no accepted view on the specific role of each area or on its network organization in humans (de Gelder and Poyo Solanas, 2021) or in monkeys (Vogels, 2022).

Another central question concerns the contribution of body perception areas to the various perceptual functions that include body perception as well as action and expression perception. Studies focusing on body perception as part of research on action and emotion recognition revealed other areas in addition to those known from category-based studies. A comparison of expressive with neutral whole body still images (de Gelder et al., 2004, 2010) and studies using video images and controlling for action category (Grèzes et al., 2007) reported the posterior superior temporal sulcus (pSTS), temporoparietal junction (TPJ), frontal cortex and parietal motor regions (Pichon et al., 2009; Peelen et al., 2007; Grèzes et al., 2007), as well as the amygdala (AMG) (de Gelder and Poyo Solanas, 2021; Poyo Solanas et al., 2020b; Pichon et al., 2012). Notably, most of the clusters found in body expression studies were also reported in studies of the action observation network (Grèzes et al., 2007; Goldberg et al., 2014; Pichon et al., 2009), emotion (de Gelder et al., 2004; Borgomaneri et al., 2015) and included subcortical areas (Poyo Solanas et al., 2020b; Utter and Basso, 2008). The relation between category-selective areas and areas that seem to be involved in perceiving various functional roles of the body is still poorly understood.

To summarize, there are now some robust findings of body category selectivity in a few different brain areas in human and monkey. This raises the question of the underlying computational processes defining their respective roles, and of the interaction of the various body selective areas in hierarchical or parallel processing streams. For example, it is unclear what the computational processes presumably taking place in each body selective area are, and whether these are best understood at the level of each separate body selective area or, alternatively, at the level of interacting body areas and network functions.

Our goal was to discover the network organization of body perception in a data-driven way rather than by investigating local areas of category selectivity for bodies (Peelen and Downing, 2005) or for body expressions (de Gelder et al., 2010). We tested the hypothesis that there might be a basic body representation network that sustains different specific domains of human body perception. To investigate human body processing at the network organization level we used ultra-high field 7T fMRI while participants viewed naturalistic dynamic videos of human and monkey faces, human and monkey bodies, and objects, as well as a scrambled version of each video as a control. Large-scale networks modulated by body processing were identified by the group independent component analysis (GICA), which has been widely used in resting-state and task-based fMRI studies (Du et al., 2017; Jarrahi et al., 2015; Jung et al., 2020). This GICA approach allowed us to separate single-voxel time courses into multiple components with maximized spatial independence. Here, the time course reflects a coherent fluctuation associated with an intrinsic network or associated with noise. Thus, by modeling the component time courses, we were able to reveal the networks modulated by our experimental conditions. Furthermore, to bring human body selectivity more narrowly in focus, we included monkey videos as the stimuli. Through the comparison with nonhuman species, it may offer insights into what exactly is coded in body selective areas and their network functions.

## RESULTS

Nineteen participants took part in the experiment. Two were excluded from further analysis due to large distortion of the functional or anatomical image. Twelve categories of videos (body/face/object * human/monkey * normal/scramble) were shown to the participants during the 7T fMRI scanning using a blocked design with six repetitions per category.

### Univariate analysis

A random-effects general linear model (GLM) with all conditions as predictors was performed to find voxel-wise (human) body preference (see Methods). To control for low-level stimulus features such as the luminance, contrast, and the amount of local motion, we computed the contrast of [human body (normal - scramble)] > [human object (normal - scramble)]. The resulting statistical map was corrected using a cluster threshold statistical procedure based on Monte Carlo simulation (initial *p* < 0.005, alpha level = 0.05, iterations = 5000). Several body selective clusters were found in the extrastriate cortex (corresponded to EBA), fusiform cortex, pSTS, TPJ, and frontal gyrus, in agreement with previous body perception studies (de Gelder and Poyo Solanas, 2021; Ross et al., 2020) (Table 1, Figure 1a). Subcortical regions including the amygdala, pulvinar, and caudate nucleus also showed body selectivity. The largest cluster corresponded to the right EBA (8355.84 mm^3^) and the highest peak t-value was found in the right amygdala (*t*(16) = 5.90, *p* < 0.001).

**Table 1.**
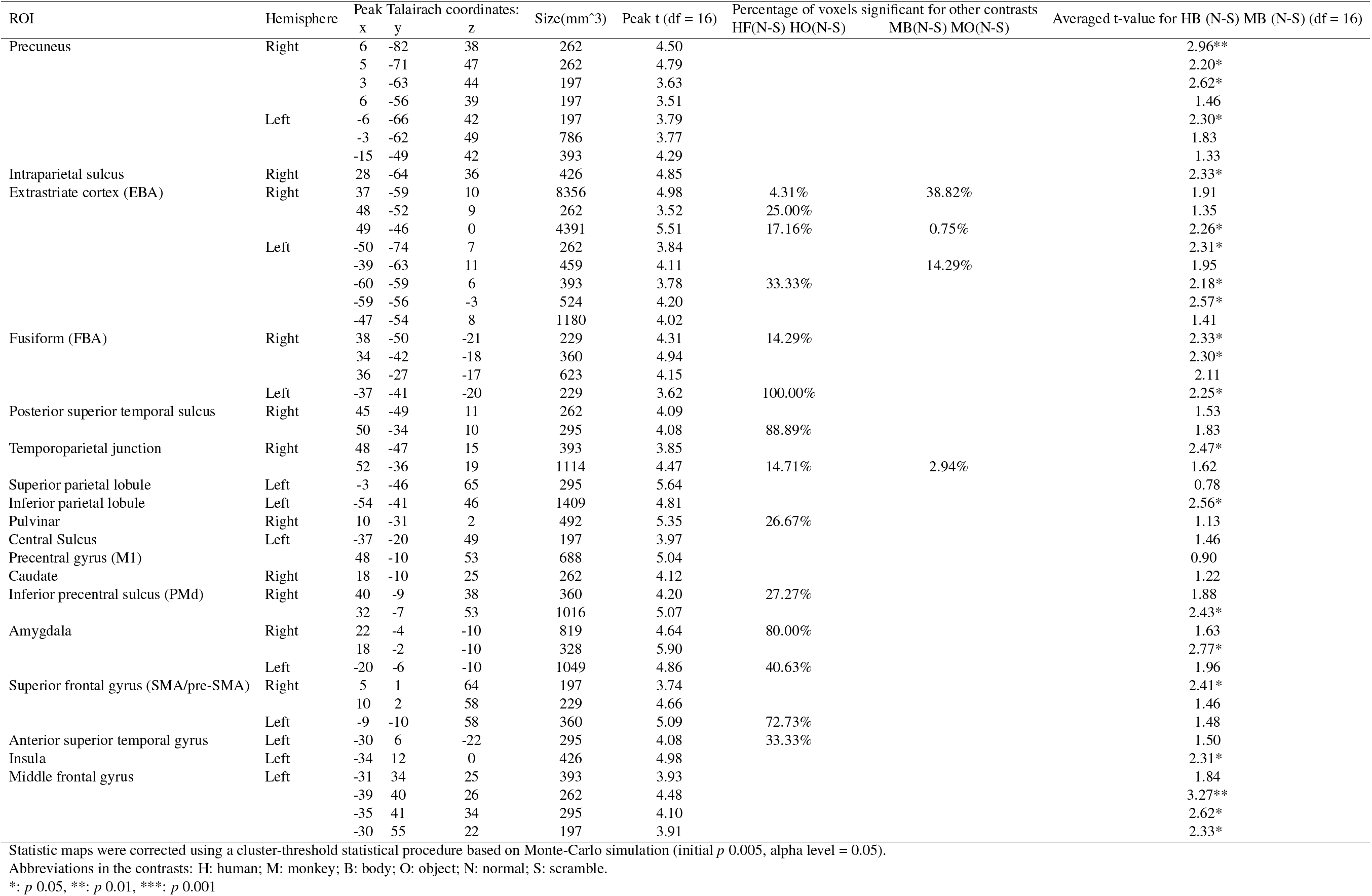
Clusters found by random-effect group GLM. Contrast: human body(normal - scramble) > human object(normal - scramble)

**Figure 1.**
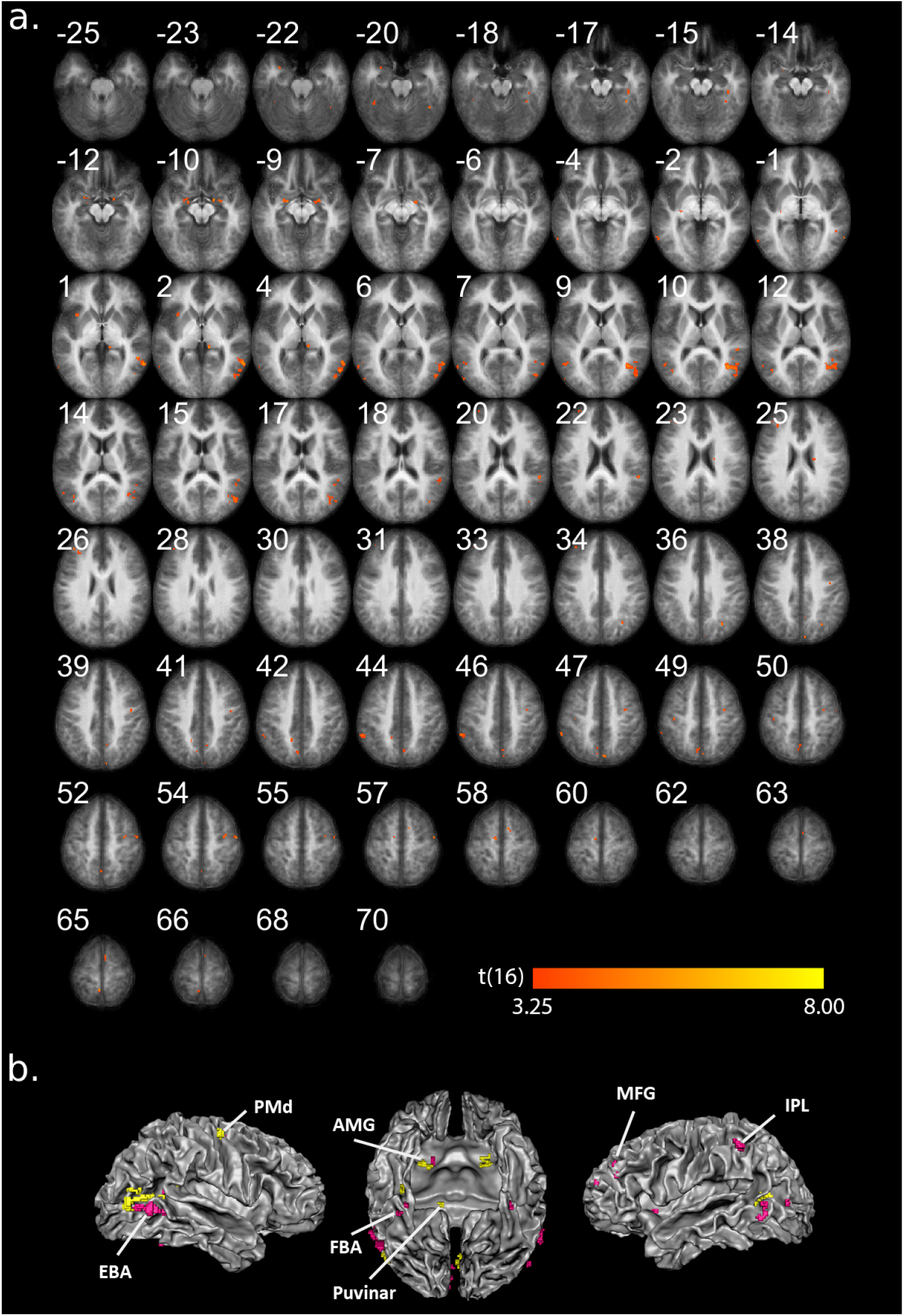
Group univariate results. **(a)**. Contrast of [HB(N-S) > HO(N-S)] (only positive values are shown). The resulting statistical map was corrected using a cluster-threshold statistical procedure based on Monte-Carlo simulation (initial *p* < 0.005, alpha level = 0.05). The number on each slice indicates the z-coordinate of Talairach space. **(b)**. The same clusters in (a) projected to the cortical mesh. ROI-level significant for contrast [HB(N-S) > MB (N-S)] are colored in pink (uncorrected *p* < 0.05, Table 2). Abbreviations in the contrasts: H: human; M: monkey; B: body; O: object; N: normal; S: scramble.

We further computed two additional low-level controlled contrasts to find a) human face selectivity by [human face(normal-scramble) > human object (normal-scramble)] and b) monkey body selectivity by [monkey body(normal-scramble) > monkey object (normal-scramble)]. After thresholding the statistical maps, overlaps were computed between the previously found human body clusters and the new contrasts. The largest overlaps were found in a) a left fusiform body cluster, where 100% of voxels were also selective to the human face, and b) a right EBA cluster, where 39% of voxels were also selective to the monkey body compared to objects (Table 1, S1 & S2).

To test the human body specificity of the body areas found above, we computed the low-level controlled contrast of [human body(normal-scramble) > monkey body (normal-scramble)] on each human body region of interest (ROI) defined above. Multiple body clusters were significantly species-selective, including EBA, fusiform, insula, middle frontal gyrus (MFG), precentral gyrus (corresponding to the dorsal premotor cortex, PMd), inferior parietal lobe (IPL) and amygdala (Figure 1b). The cluster showing the highest human specificity was found in the MFG (*t*(16) = 3.27, *p* = 0.005, Table 1).

### Independent Component Analysis

To study the network organization related to body perception, we applied a data-driven approach with group independent component analysis (GICA). Seventy-five independent components (ICs) were extracted from the preprocessed data (see Methods). A systematic pipeline was applied to exclude noise components and to find category-modulated networks. Five components were first removed due to an ICASSO Iq value lower than 0.8 (Himberg et al., 2004). The positive and negative parts of the remaining ICs were further divided into different IC sets, and the sign of the time courses and spatial maps of the negative ICs were flipped. Of the resulting 140 ICs, 16 positive ICs and 28 flipped ICs were identified as noise and were excluded due to white matter (WM) / cerebrospinal fluid (CSF) overlap larger than 10%. Task relevance was modeled for each reconstructed IC time course using a GLM with the same design matrix as in the univariate analysis. Here, we assumed a positive hemodynamic response function (HRF) response for the cortical network time courses, thus the ICs / flipped ICs with a negative mean beta across all conditions were excluded from further analysis. Finally, 19 positive ICs and 31 flipped ICs were used in further analyses.

To investigate condition-specific modulations within these ICs, several contrast analyses were conducted with the estimated betas from the IC time courses. For the first contrast of [normal human body > normal human object], we found only one network showing significant selectivity for human bodies after multiple comparison corrections (IC42, Figure 2a, *t*(16) = 3.97, Benjamini-Hochberg False Discovery Rate corrected *q* < 0.05, right-tailed). The network (referred to as the rSTS network for abbreviation) covered right-lateralized regions including EBA, fusiform, STS, TPJ, IPL, MFG, precentral gyrus (PrCG), inferior frontal gyrus (IFG) and pulvinar, as well as bilateral clusters around amygdala, insula and supramarginal gyrus (SMG). Further inspection of the estimated betas revealed a significant preference of this network for human faces over monkey faces (*t*(16) = 2.40, *p* = 0.029, two-tailed) and for human bodies over monkey bodies (*t*(16) = 2.92, *p* = 0.010, two-tailed) (Figure 2c). Further inspection of the beta plot revealed a structural response profile where the highest response was found for the human face, then the human body and the monkey face, and the monkey body came to the last (Figure 2c). However, the response difference was not significant between human body and human face conditions (*t*(16) = 1.77, *p* = 0.096, two-tailed)

**Figure 2.**
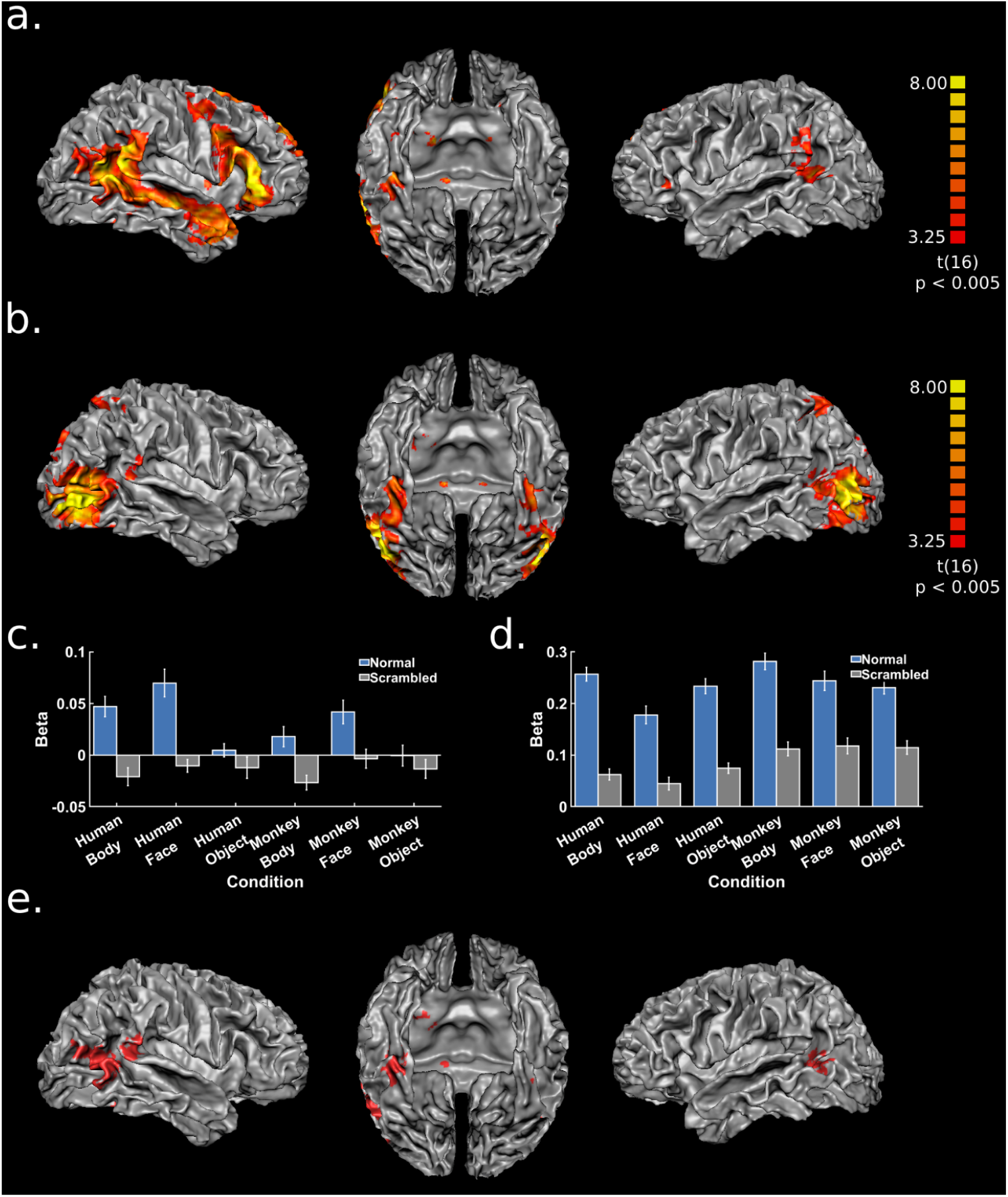
Networks extracted by group-ICA. The individual IC maps were z-transformed and averaged across all runs for each participant. A group t-test against zero was computed using the z-scored maps of each subject. The resulting statistical map was corrected using a cluster-threshold statistical procedure based on Monte-Carlo simulation (initial *p* < 0.005, alpha level = 0.05). **(a) & (c)**. rSTS network and its beta plot. **(b) & (d)**. LOC network and its beta plot. **(e)**. The overlap between the two networks.

For the second contrast analysis, we controlled for low level features. Using the contrast of [human body (normal - scramble) - human object (normal - scramble)]), in addition to the rSTS network (*t*(16) = 2.93, corrected *q* < 0.05, right tailed), another IC also showed human body selectivity (IC04, Figure 2b, *t*(16) = 3.29, corrected *q* < 0.05, right-tailed). The spatial map of this component revealed a lateral occipital cortex dominant network (referred to as the LOC network for abbreviation), which also included bilateral fusiform, superior parietal lobe (SPL), pSTS/TPJ, pulvinar and amygdala. However, no human specificity was found either by the contrast of [human body (normal - scramble)] > [monkey body (normal - scramble)] (*t*(16) = 1.98, *p* = 0.065, two-tailed), or by the contrast of [human face (normal - scramble)] > [monkey face (normal - scramble)] (*t*(16) = 0.51, *p* = 0.615, two-tailed) (Figure 2d). The contrast of [human body (normal - scramble)] > [human face (normal - scramble)] revealed a significant preference for human body over human face (*t*(16) = 4.12, *p* < 0.001, two-tailed). Overlap between the rSTS network and the LOC network was found around the temporo-occipital region, covering the clusters of EBA, fusiform, pSTS, TPJ as well as pulvinar and amygdala (Figure 2e), which were also found by univariate analyses.

To further investigate condition-specific modulations on the node connectivity of the above-mentioned networks, we repeated the same ICA procedure after regressing out the activity of one category from the time courses and we compared the condition-omitted spatial maps and the original one for the same network. With this comparison, the condition dependence of the nodes can be then identified as decreased network weights after the omission. As a result, significant drops in IC weight were detected in EBA, pSTS/TPJ, PrCG (corresponding to PMd/PMv) and IFG in the rSTS network after the normal human body blocks were omitted (Table 2, Figure 3). Both the largest cluster and the peak t-value were found in IFG (largest V = 14647.30 mm3; highest peak *t*(16) = 7.60, *p* < 0.001). For the LOC network, the connectivity weight drops were observed mainly around bilateral EBA (Table 2, Figure 3), with the largest cluster and peak t-value found in right EBA (largest V = 6815.74 mm3; highest peak *t*(16) = 6.70, *p* < 0.001).

**Table 2.**
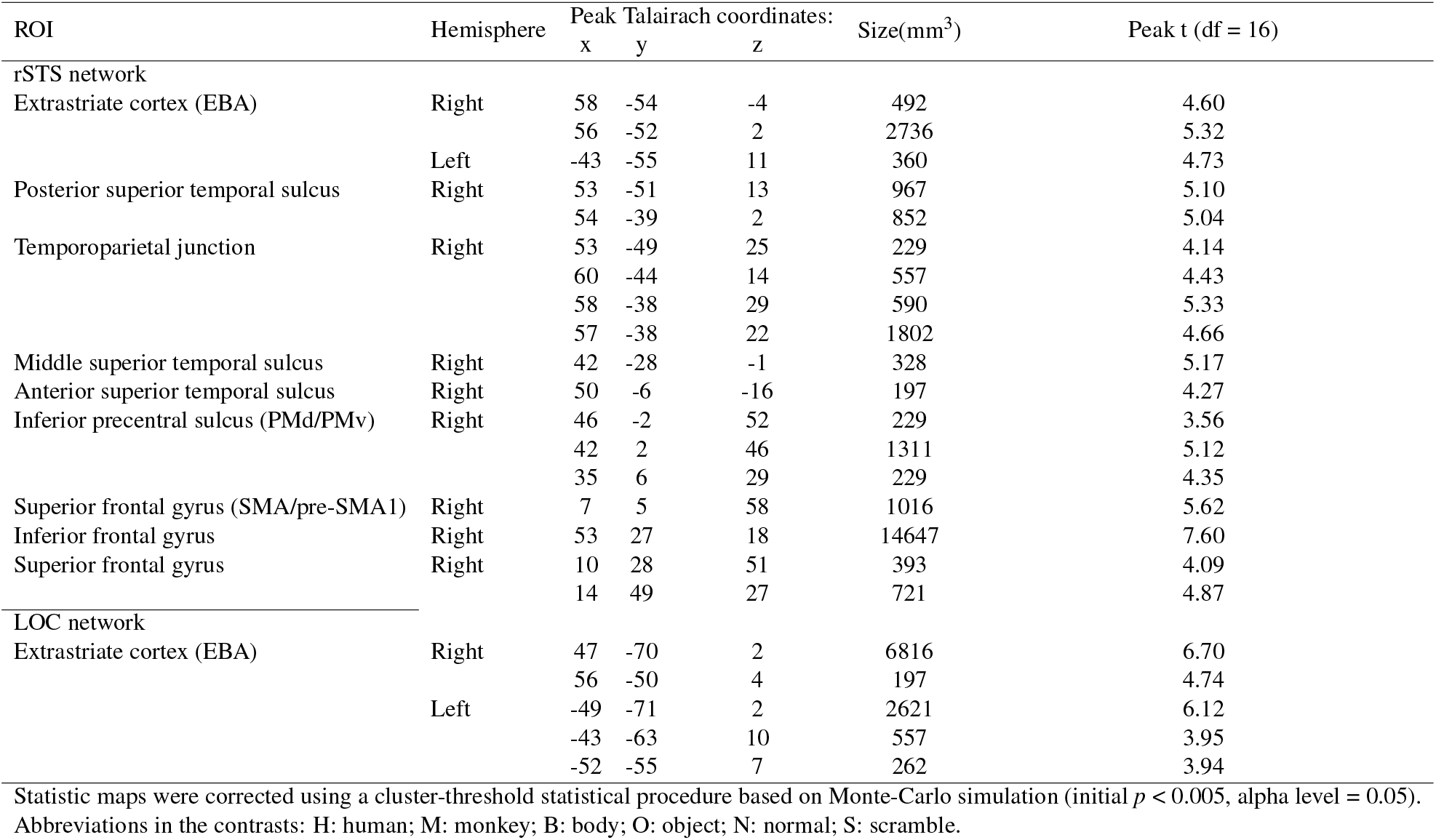
Clusters found by original ICA - HB-omitted ICA

**Figure 3.**
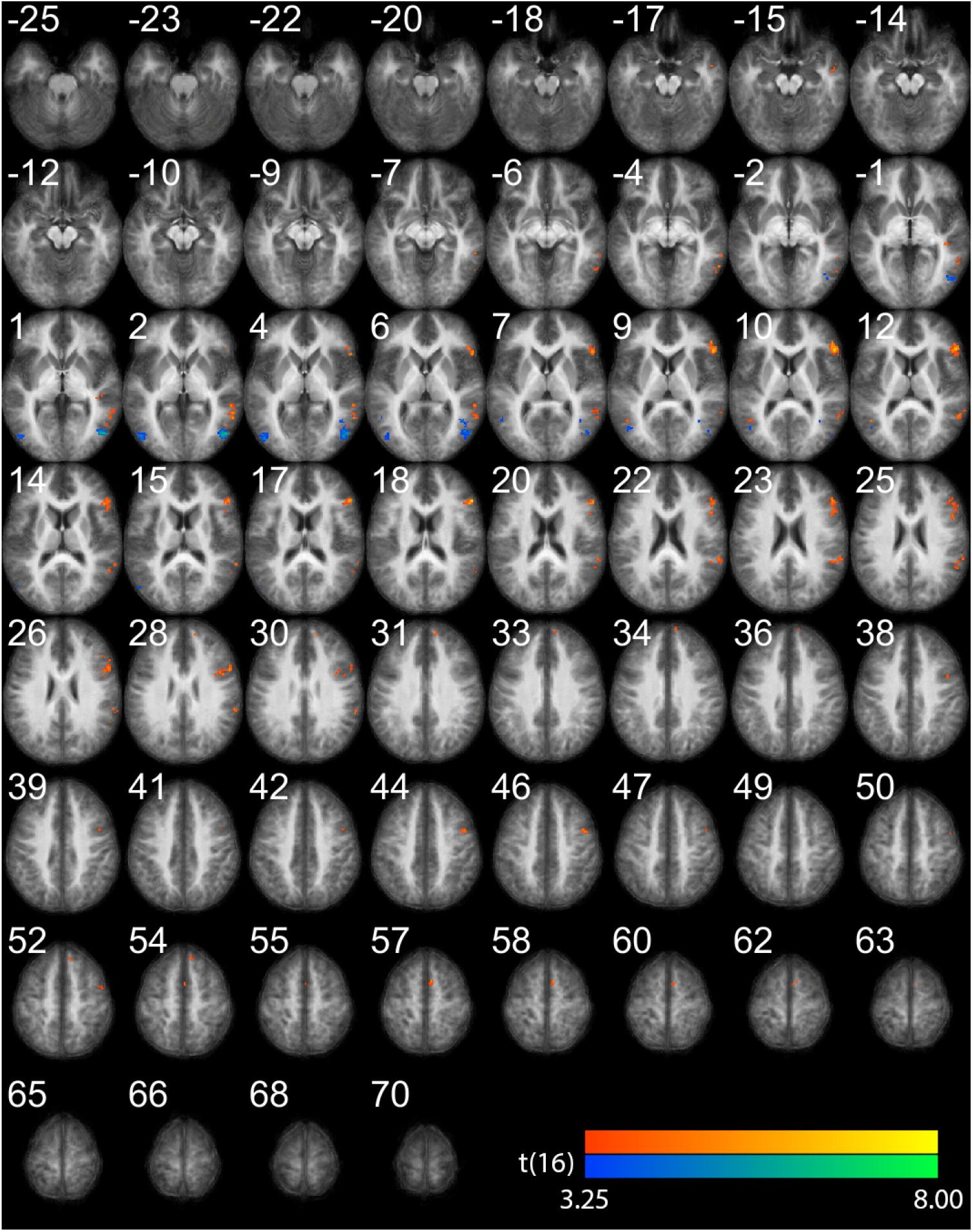
Connectivity drops calculated by original ICA – HB-omitted ICA for the two networks. The group statistical map was corrected using a cluster-threshold statistical procedure based on Monte-Carlo simulation (initial *p* < 0.005, alpha level = 0.05) and masked by thresholded networks in Figure 2a&b separately. Red clusters indicate significant connectivity drops for rSTS network, and blue clusters indicate drops in LOC network.

In addition to defining the body nodes, we reconstructed the networks separately after regressing out the human face condition and the monkey body condition. Within the defined body nodes, we first searched for the voxels showing significant connectivity decrease for human-face-regressed and monkey-body-regressed maps. For the rSTS network (Figure 4a), the human face dependence was found in the right pSTS, TPJ, PMd and IFG body nodes (uncorrected *p* < 0.05, Figure 4b). Monkey body dependence was only found around the right EBA and pSTS body node (uncorrected *p* < 0.05, Figure 4c). Next, to find voxels with unique dependence on the human body, we conducted a conjunction analysis with the contrast of [decrease(human body) > decrease(human face)] and [decrease(human body) > decrease(monkey body)] within the body nodes. As a result, significant voxels were found in the bilateral EBA, right TPJ, PMv, SMA, SFG, and IFG body nodes (uncorrected *p* < 0.05, Figure 4d). For the LOC network, voxels with monkey-body or human-face-dependent voxels were found in bilateral EBA nodes, while the human-body-specific voxels were mainly found in the left EBA node.

**Figure 4.**
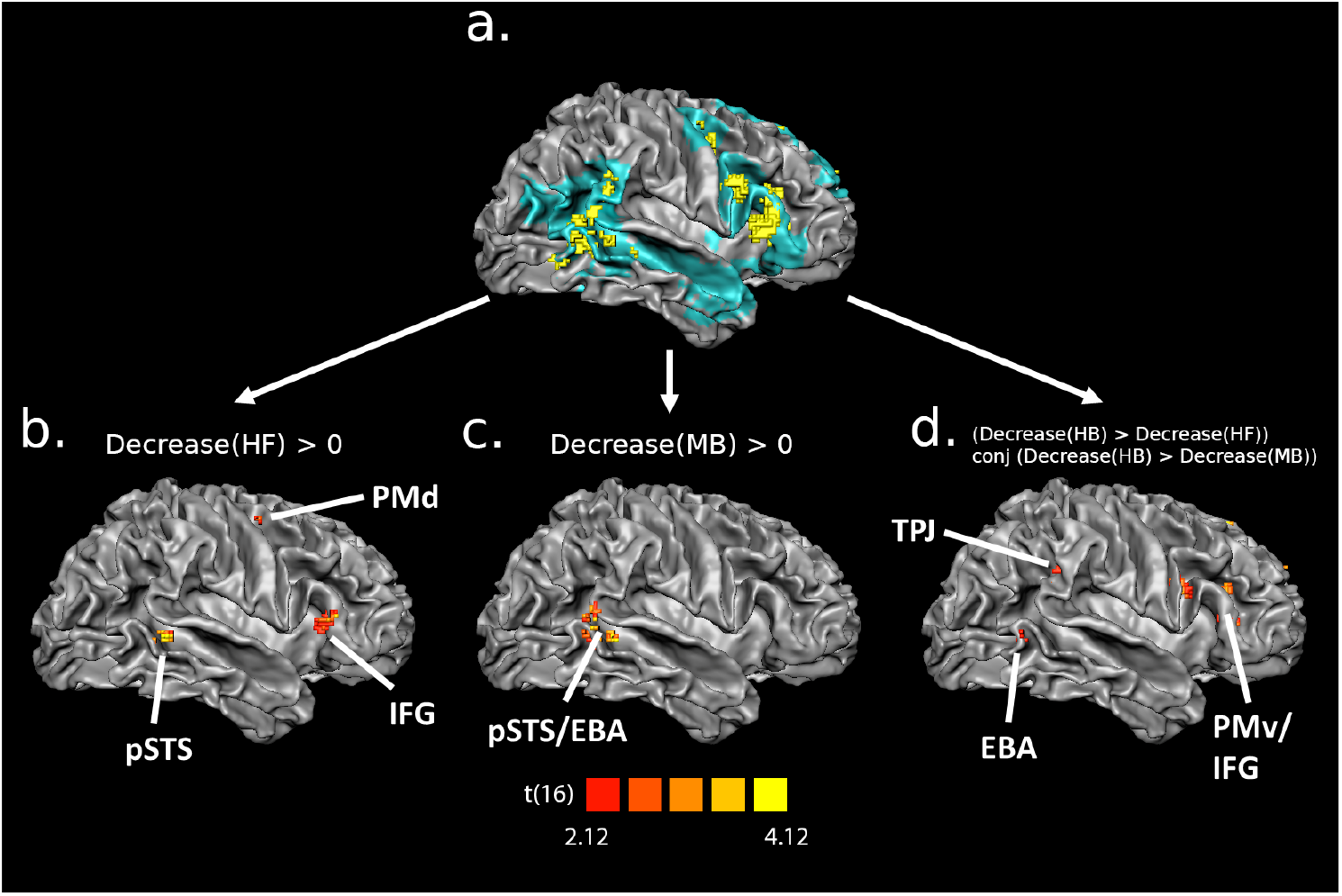
Dependence properties revealed by the weight decreases for the rSTS body nodes. **(a)**. The rSTS nodes in Figure 3 projected to cortical mesh with blue shadows indicating the network coverage. **(b)**. Node voxels showing human face dependence. **(c)**. Node voxels showing monkey body dependence. **(d)**. Node voxels showing human-specific body dependence. Abbreviations in the contrasts: H: human; M: monkey; B: body; F: Face.

## DISCUSSION

Using dynamic multispecies stimuli, 7T fMRI scanning and data-driven methods we investigated body selective areas and their species specificity and category selectivity and focused on the network organization of body processing. Our analyses discovered two large-scale networks specifically modulated by human body videos, the LOC network and the rSTS network. As the study used novel video materials, we first briefly discuss these new findings on body selectivity and species specificity in the light of the literature and then address the main finding of the network connectivity, and we indicate the novelty of our network findings in contrast with earlier proposals based on a priori higher-order stimulus categorization. Finally, we propose an interpretation of the possible functions of the two body-modulated networks.

### Multiple areas of body selectivity

Our univariate results provide the first complete picture based on ultra-high field scanning of areas involved in dynamic body processing that are specific to the human body. First, concerning EBA and FBA, our results are consistent with those of previous studies using videos (de Gelder and Poyo Solanas, 2021). Our novel result here is that a subset of EBA and fusiform clusters showed higher responses for human bodies than for monkey bodies. A possible basis for human body specific coding may be that these areas compute features that are more characteristic of human body movements, for example, because they abide by biomechanical constraints of human body posture and motion. A related basis for human-specificity also at the feature level may be that the coding in these two areas is partly driven by expression perception. For example, the features that deliver some affective information embedded in human body expressions (Poyo Solanas et al., 2020b,a) may be absent in monkey bodies.

Two other areas, pSTS and TPJ are mostly known for their involvement in dynamic face processing (Patel et al., 2019). Still, they have appeared already since the first studies on body perception (de Gelder et al., 2004) as well as in later ones (Pitcher et al., 2019; Kret et al., 2011; Grèzes et al., 2007). We further detailed at this by showing that in some parts of this pSTS/TPJ cluster there is human-body-specific coding. This may provide a conceptual basis for previous findings on the biological motion of faces and bodies (Patel et al., 2019; Polosecki et al., 2013; Yovel and O’Toole, 2016). Other recent studies have proposed that these two regions may be related to the predictive coding of biomechanical movements (Geng and Vossel, 2013; Koster-Hale and Saxe, 2013). pSTS/TPJ is involved in the generation of model-based predictions of biomechanical trajectories of moving face or body parts while also updating the models according to the new incoming information (Patel et al., 2019; Geng and Vossel, 2013; Koster-Hale and Saxe, 2013).

We also found several human body-selective clusters in the frontoparietal and subcortical regions. Frontoparietal areas include SPL, intraparietal sulcus (IPS), as well as PMd and belong to the dorsal frontoparietal network (dFPN), which may be involved in the dynamic representation of the kinematic properties of movement plans (Ptak et al., 2017). Finally, subcortical clusters were found in the pulvinar and amygdala. The amygdala has been reported to detect behaviorally relevant stimuli and has also been previously observed for body images (Hadjikhani and De Gelder, 2003) and videos (Grèzes et al., 2007; Pichon et al., 2009).

### Network-based body representation

Our ICA analysis discovered two networks that showed significant modulation by body stimuli and had very different response profiles for the other categories. In both networks, we found nodes that had their connectivity significantly influenced by bodies. Some of these node-level modulations also showed human specificity, especially in the rSTS network.

#### LOC network

The LOC network mainly consisted of a large cluster in the lateral occipital cortex and the fusiform cortex, covering most of the previously defined category-selective areas (Grill-Spector and Sayres, 2008). The classical view of category-selective areas is that these areas compute the entry-level representation of the preferred category and that these category computations are not dependent on low level features. But the current understanding of the relationship between low-level features (contrast edges, local motion, luminance, differences in spatial frequency) and high-level category-defining representation is limited (Long et al., 2018; de Gelder and Poyo Solanas, 2021). In this respect, it is interesting to see that the body selectivity of this network emerged when taking the respective scrambled control conditions into account. Thus, the LOC network may be selective for specific properties of the body videos (Grill-Spector and Weiner, 2014) and this selectivity may be partly based on midlevel features like human body specific movement or postural characteristics over time (Poyo Solanas et al., 2020b,a).

#### rSTS network

The rSTS network showed a right hemisphere-dominant coverage including EBA, FBA, STS, PMd/PMv and IFG. Other nodes covered by the rSTS network, such as the premotor cortex, medial prefrontal cortex, TPJ and amygdala, have also been frequently related to social cognition (Saxe and Kanwisher, 2003; Schurz et al., 2014; Van Overwalle, 2009; Young et al., 2010; Patel et al., 2019; Alcalá-López et al., 2018). Most notably, this network showed the highest response for human faces and human bodies, followed by monkey faces, and lastly monkey bodies (Figure 2c). While the contrast was not significant between human bodies and faces, significantly higher responses were found for human videos than for the monkey ones. Thus, the rSTS network may involved the processing of human-specific social information.

#### Node-level body modulation within networks

In addition to finding body modulations at the network level, we were interested in identifying the nodes within each network that were involved in body processing compared to the other stimulus conditions. Using condition-omitted ICA, we first found body modulations of node connectivity only in the bilateral posterior EBA in the LOC network. Similarly, EBA nodes were also body-modulated in the rSTS network, which overlapped with the anterior EBA cluster found to be human-specific in our univariate analysis. It should be noted that while the anterior EBA was covered by both the LOC and rSTS networks, the posterior EBA was only covered by the LOC network. This result suggests that the posterior and anterior EBA may be involved in different information flow during body processing. This could be presumably related to the different contributions or different computations of the EBA subparts in each network. This proposal is consistent with the notion that EBA is a complex area covering three heterogeneous regions surrounding the human motion-selective complex (hMT+) (Weiner and Grill-Spector, 2011).

In the rSTS network, more body-modulated involvement was also besides EBA, in TPJ, premotor cortex, frontal gyrus, and the clusters along STS. A notable property of the current rSTS network is its right lateralization, which was previously only found in studies on face processing(De Winter et al., 2015; Sato et al., 2019; Yokoyama et al., 2021). Interestingly, other studies suggested an opposite view of the lateralized social network, with the left hemisphere related to the detailed evaluation of social signals and the right hemisphere to rapid automatic detection of the high valence stimuli(Alcalá-López et al., 2018). Such contrasting views may indicate that, between the low-level visual features and the full extraction of semantic information, there are intermediate stages during the processing, especially of the affective social signal.

#### The subnetwork for human-specific body processing

To consolidate the evidence in favor of the human body specificity of the nodes detected above, we further searched for the voxels with distinct or shared dependence for human bodies compared to the human face and monkey body. The result showed that among the rSTS body nodes, voxels within the EBA, TPJ, PMv, SMA, SFG and IFG nodes showed significantly larger connectivity decreases for the human-body-regressed network than for the human-face- or monkey-body-regressed ones. This result suggested a subnetwork for human-specific body processing.

Moreover, human-face dependent voxels were also found in body nodes around pSTS, PMd, and IFG, suggesting that the common features between body and face, such as biological motion and social information, may be processed here. The IFG node is at the intersection of the human-body-specific and the body-face-shared subnetworks and may be crucial for understanding human-specific body and social information. The IFG has been associated with multiple cognitive functions, including attention, social cognition, and motor inhibition (Hartwigsen et al., 2019). However, in the context of body perception and connectivity, one crucial property of right IFG is its connection to TPJ. The right IFG and the right TPJ are anatomically connected by the third tract of the superior longitudinal fasciculus (SLF III), which was reported to be highly anatomically asymmetric (Wang et al., 2016). Moreover, lesions in SLF III are often related to dysfunctions in embodiment that can cause patients to misidentify others’ limbs as their own (Errante et al., 2022). Thus, the cooccurrence of the IFG and the TPJ nodes in the human-body-specific subnetwork may suggest a stronger involvement of embodiment when viewing human body videos.

The right IFG was also reported to be selective to biological motion and dynamic bodies and the connectivity between IFG and pSTS is sensitive to biological motion (Saygin et al., 2004; Jung et al., 2009; Ross et al., 2020; Sokolov et al., 2018). Other studies linked the right IFG to a predictive coding leading to the detection of the mismatch between the actions and their context (Wurm and Schubotz, 2012; Hrkać et al., 2014; Urgen and Saygin, 2020). Furthermore, both TPJ and IFG were reported to be involved in the model-based prediction and inferences about the state of the agent from the actions(Koster-Hale and Saxe, 2013). This prediction perspective is also compatible with the current results, especially for the voxels showing shared dependence on the human body and face.

In addition to the IFG node, the pSTS node in the rSTS network also showed a notable property. While multiple body nodes including pSTS were found with human-face dependence, the pSTS node (and a small proportion of EBA) was the only one with monkey body dependence. This suggests that on one hand, pSTS may serve as a starting point to integrate the general features of body and face with no species selectivity. But, for the monkey body, such information may be further gated before being sent to the other nodes. This explanation is compatible with the proposal of Patel et al. (2019), who suggested the pSTS sends inputs to the TPJ and participates in a larger network. However, how the nonsocial or non-human information is filtered out is still a question for future studies.

#### Correspondence and intersection between the two networks

An interesting question concerns the communication between the two networks. Thus, we further inspected the overlaps between the LOC and rSTS networks, aiming to find a potential bridge linking the lower- and higher-level processing of body stimuli. Besides the regions of the EBA and FBA,the most notable cortical intersections of the two networks were found around pSTS/TPJ, which is again compatible with the notion of pSTS/TPJ as a middle-station between networks mentioned above. The connection between lower- and higher-level information can also be found in the pulvinar region, which was found as a main subcortical intersection between our two networks. As mentioned above, the ventral part of the pulvinar is sensitive to low-level temporal structures, while the dorsal part is selective to more integrated information (Arcaro et al., 2018; Hasson et al., 2008, 2015). Consistent with this, only the ventral pulvinar was involved in the LOC network, while both the ventral and dorsal parts were found in the rSTS network. In conclusion, pSTS/TPJ and pulvinar may play an important role during information exchanges between the lower-level feature system and the higher-level social information system.

### Relation between category, action and emotion perception and the social brain networks

The present rSTS network was found using a data-driven approach with dynamic body stimuli and using ultra-high-field fMRI. Previous studies each defined somewhat similar networks using different stimuli and tasks and other network proposals were based on meta-analyses or used data from the human connectome project (e.g., Alcalá-López et al. 2018). The first network is the action observation network (AON, Caspers et al., 2010), with similar nodes around EBA, IFG, and PM. However, compared to the AON, our rSTS network showed a highly right-lateralized distribution that covered a large area in the right STS, which is missing in the AON. Another recent proposal on the third visual pathway stressed the role of STS in processing social information, however, this misses the links between the STS route and the other cortical regions (Haak and Beckmann, 2018). Another network proposal that has the best compatibility with our network results is a TPJ/pSTS-centered social cognition network (Patel et al., 2019). In this network, the TPJ/pSTS served as a hub receiving the input from the lower visual regions while sending integrated information to a social cognition network. Moreover, the study suggested that the third pathway of STS may serve as an input to the hub of TPJ/pSTS, thus also explaining the involvement of the large STS in our network. These findings are in line with the view that the pSTS/TPJ may serve as a hub node for integrating different functional networks (Patel et al., 2019). Our results now add that this hub function may to an important extent be based on receiving inputs from EBA/FBA.

Secondly, some nodes of the present rSTS network, such as the PMd and IPL, have been reported in studies on the role of mirror neurons in action perception (Yokoyama et al., 2021). Mirror neuron theorists argue that motor area activity seen in action perception studies is evidence for resonance and that this plays an active role in perception (as typically also argued by embodied simulation theories) (Gallese and Sinigaglia, 2011). A recent study directly tested the perception versus motor resonance hypotheses (Borgomaneri et al., 2015) and found that the early stages (150ms) of M1 reactivity corresponded to visual perception while the later stage (300ms) involved motor resonance or embodiment using mirror mechanisms.

## CONCLUSION

Our results show that the human body has a special status for human observers and may play a foundational role in more specific functional networks of social perception. This special status has different correlates. The finding that the human body network includes areas beyond the classical ventral stream one, which are associated with action and emotion perception suggests that body selectivity processes, hitherto associated with category selectivity, are tightly interwoven with processing the functional properties of bodies. In a departure from classic models of object perception, seeing human body images, specifically dynamic ones, triggers a functional network-based representation, rather than a neutral, context-free category representation more directly than objects do. Next, this functional network representation may be model-based, driven by an internal model in the perceiver of the whole body that may be spread over multiple processes or based on network connectivity between different brain areas.

## MATERIALS AND METHODS

### Participants

Nineteen healthy participants (mean age = 24.58; age range = 19-30; 6 males, all right handed) took part in the experiment. All participants had a normal or corrected-to-normal vision and no medical history of any psychiatric or neurological disorders. All participants provided informed written consent before the start of the experiment and received a monetary reward (vouchers) or course credits for their participation. The experiment was approved by the Ethical Committee at Maastricht University and was performed in accordance with the Declaration of Helsinki.

### Stimuli

The materials used in this experiment consisted of 1-second-long grayscale videos of bodies, faces, and objects edited from original human and monkey recordings. The body and face videos depicted either a human or a monkey performing naturalistic full-body or facial movements. Object stimuli consisted of two sets of moving artificial objects with the aspect ratio matched to either human bodies or monkey bodies. The size of the stimuli was 3.5*3.5 degrees of visual angle for human faces, 3.5*7.5 degrees for human bodies and objects, and 6*6 degrees for monkey faces, bodies and objects. The human videos were selected from the set originally developed in Kret et al. (2011), in which all actors were dressed in black and performed natural full body / face expressions against a greenscreen background. The expressions contained anger, fear, happiness, as well as neutral expressions such as pulling nose or coughing. The monkey videos were taken from footage of rhesus monkeys from the KULeuven monkey colony and also from a published comparative study of facial expressions Zhu et al. (2013). The body videos included grasping, picking, turning, walking, threatening, throwing, wiping, and initiating jumping, while the face videos included chewing, lip-smacking, fear grin, and threat. For human and monkey videos, a variety of both emotional and neutral poses were included, and the face information within each body video was removed by applying Gaussian blurring.

After removing the original background, the videos were cut to 1s duration (60 frames/s) and overlaid on a full-screen dynamic white noise background spanning 17.23*10.38 degrees of visual angle. The background consisted of small squares of 3 by 3 pixels of which the gray level was randomly sampled from a uniform distribution at a rate of 30 Hertz. To directly control for low-level feature differences among the three categories (bodies, faces and objects), we included mosaic-scrambled videos as an additional set of stimulus conditions. The mosaic scrambled stimuli destroyed the whole shape and global motion of the dynamic bodies, faces, and objects, but had identical local motion (within 14 pixels wide squares), luminance, contrast, and non-background area as the original movies. This resulted in a total of twelve experimental conditions (human/monkey * body/face/object * normal/scrambled). There were ten different stimuli per condition, which resulted in 120 unique videos.

### Experimental design

During the experiment, stimuli were presented following a block-design paradigm. For each block, ten videos of the same experimental condition were presented once for 1000 ms in random order with an inter-stimulus-interval (ITI) of 500-ms consisting of a uniform gray canvas. Two blocks per condition were randomly presented within each run. Between blocks, there was a jittered interval of 11s where a blank canvas was presented. For each participant, we collected three experimental runs, resulting in six repetitions per condition. At the beginning and the end of each run, a white noise block was presented with only the dynamic noise background but no actual stimulus (ten videos of 1-second with an ITI of 500-ms). Ultimately, for each run we collected 735 functional volumes resulting in approximately 12 minutes of scanning time.

During the experiment, participants were instructed to keep fixation on a cross presented at the center of the screen throughout the whole run. Participants’ attention was controlled by adding two catch blocks in each run, in which the fixation cross changed its shape to a circle during a random trial. The participants were asked to press a button with the right index finger when detecting the fixation shape change. The category of each catch block was randomly chosen from the twelve experimental conditions, and all of the catch blocks were removed from further data analysis to rule out response-related confounds.

The experiment was programmed using the Psychtoolbox (https://www.psychtoolbox.net) implemented in Matlab 2018b (https://www.mathworks.com). Stimuli were projected onto a screen at the end of the scanner bore with a Panasonic PT-EZ57OEL projector (screen size = 30 * 18 cm, resolution = 1920 * 1200 pixel). Participants viewed the stimuli through a mirror attached to the head coil (screen-to-eye distance = 99 cm, visual angle = 17.23 * 10.38 degrees). The whole experiment lasted for 40 minutes. The same participants underwent another round of scanning for a different experiment which is not reported here.

### fMRI data acquisition

All images were acquired with a 7T MAGNETOM scanner at the Maastricht Brain Imaging Centre (MBIC) of Maastricht University, the Netherlands. Functional images were collected using the T2*-weighted multi-band accelerated EPI 2D BOLD sequence (TR/TE = 1000/20 ms, multiband acceleration factor = 3, in-plane isotropic resolution = 1.6mm, number of slices per volume = 68, matrix size = 1152 * 1152, volume number = 735). T1-weighted anatomical images were obtained using the 3D-MP2RAGE sequence (TR/TE = 5000/2.47 ms, Inverse time TI1/I2 = 900/2750 ms, flip angle FA1/FA2 = 5/3°, in-plane isotropic resolution = 0.7mm, matrix size = 320 * 320, slice number = 240). Physiological parameters were recorded via pulse oximetry on the index finger of the left hand and with a respiratory belt.

### fMRI image preprocessing

Anatomical and functional images were preprocessed using the Brainvoyager 22 (Goebel, 2012) and the Neuroelf toolbox in Matlab (https://neuroelf.net/). For anatomical images, brain extraction was conducted with INV2 images to correct for MP2RAGE background noise. For functional images, the preprocessing steps included EPI distortion correction (Breman et al., 2020), slice scan time correction, 3D head-motion correction, and high-pass temporal filtering (GLM with Fourier basis set of 3 cycles, including linear trend). Coregistration was first conducted between the anatomical image and its most adjacent functional run using a boundary-based registration (BBR) algorithm (Greve and Fischl, 2009), and all the other functional runs were coregistered to the aligned run. Individual images were normalized to Talairach space (Collins et al., 1994) with 3 mm Gaussian spatial smoothing. Trilinear/sinc interpolation was used in the motion correction step, and sinc interpolation was used in all of the other steps.

Physiological parameters were collected as the confounds of functional imaging data. The physiological data were preprocessed using the RETROspective Image CORrection (RETROICOR; Glover et al., 2000; Harvey et al., 2008) pipeline, which uses Fourier expansions of different orders for the phase of cardiac pulsation (3rd order), respiration (4th order) and cardio-respiratory interaction (1st order). 18 physiological confounds were finally created for each participant.

For visualization, we created a cortical mesh from a single subject in Talairach space. The subject anatomical image first underwent a fine-tuned deep-learning-based segmentation implanted in Brainvoyager. The resulting gray/white matter labeling image was then aligned to the group-averaged anatomical image with SyN algorism using the toolbox of Advanced Normalization Tools (ANTs; Avants et al., 2022). The group cortical mesh was finally created from the aligned labeling image.

### Univariate analysis

A random-effects general linear model was performed to find the voxel-wise categorical preference. In the design matrix, each condition predictor was modeled as a boxcar function with the same duration of the block and convolved with the canonical hemodynamic response function (HRF). Physiological and motion confounds were added as nuisance repressors.

Body selective areas were defined by the contrast analysis of [human body (normal - scrambled) > human object (normal - scrambled)]. The term of (normal - scramble) aimed to rule out influences from low-level stimulus features. The resulting statistical map was corrected using a cluster-threshold statistical procedure based on the Monte-Carlo simulation (initial p < 0.005, alpha level = 0.05, iteration = 5000).

Besides the body contrasts, we calculated two additional low-level controlled contrasts for human face selectivity [human face (normal - scrambled) > human object (normal - scrambled)] and cross-species body selectivity [monkey body (normal - scrambled) > monkey object (normal - scrambled)]. The statistical maps were thresholded in the same manner as for the body contrasts, and the overlaps were computed for each previous body cluster and the new contrast, resulting in a proportion of voxels showing other selectivity in each body cluster.

To test the species-selectivity of the body clusters, we calculated the low-level controlled contrast of [human body (normal - scramble) > monkey body(normal - scramble)] on each body ROI. For each body cluster detected above, the t-values were averaged across all voxels, reflecting the significance of species-selectivity for bodies at a cluster level.

### Group independent component analysis (ICA)

#### ICA source data

Before performing the group-ICA, physiological and motion confounds were regressed out from the preprocessed functional images. To remove motor-related modulations, the BOLD responses for the catch blocks were removed using the finite impulse response (FIR) model. Twenty-five predictors covering 25 seconds after the block onset for each catch block were modeled and were then regressed out from the time courses using a GLM. The resulting time courses were then transformed into percentages of signal change to enhance the ICA stability (Allen et al., 2011).

#### Network extraction

Seventy-five spatial independent components (ICs) were extracted using the Infomax algorithm implemented in the Group ICA of fMRI Toolbox (GIFT, Calhoun et al., 2001). According to previous literature, the model of 75 components is able to cover the known anatomical and functional segmentations (Allen et al., 2011). Individual ICs were back-reconstructed using the GIG-ICA algorithm from the aggregated group ICs (Du and Fan, 2013). The stability of group ICA was assessed by the ICASSO module implemented in the GIFT, which repeated the Infomax decomposition for 20 times and resulted in an index of stability (*Iq*) for each IC (Himberg et al., 2004). To visualize the spatial map of the IC networks, the individual IC maps were normalized to z-scores and averaged across all runs for each participant. A group t-test against zero was computed using the z-scored maps of each subject and corrected using a cluster-threshold statistical procedure based on Monte-Carlo simulation (initial *p* < 0.005, alpha level = 0.05, iteration = 5000).

#### Body modulation detection

After extraction and back-construction, the individual ICs were analyzed with a data-driven approach. A systematic pipeline was applied to exclude noise components and to find category-modulated networks. ICs with an ICASSO Iq < 0.8 were first marked as unstable components and removed (Allen et al., 2011). Next, since the sign of the IC time course was arbitrary, we analyzed the positive and negative parts of each IC separately as different networks with the time courses and spatial maps flipped for the negative ones. We further labeled the white matter (WM) and cerebrospinal fluid (CSF) voxels of each thresholded IC map using customized WM / CSF masks. ICs with more than 10% WM or CSF voxels were removed as noise signals such as head motions and venous artifacts. Task relevance was modeled for each reconstructed subject-level IC time courses using a GLM with the same design matrix as in the univariate analysis and was conducted for each participant and each run separately. Such a modeling strategy was commonly used to detect the task modulations on IC networks (Beldzik et al., 2013; Jarrahi et al., 2015; Jung et al., 2020). We also assumed a positive HRF response for the cortical network time courses, thus the ICs / flipped ICs with a negative mean beta across all conditions were excluded from further analysis. Finally, we conducted a contrast analysis to find the body-selective networks. The estimated betas were first averaged across all runs for each participant and were then used to calculate the contrast of [normal human body - normal human object] and [human body (normal - scramble) - human object (normal - scramble)]. Right-tailed t-tests and Benjamini-Hochberg multiple comparison corrections were conducted at the group level to find significant body sensitivity.

#### Condition-omitted ICA

To study the body modulations on node connectivity within networks, we developed a condition-omitted ICA strategy. A human body-omitted dataset was created from the original ICA source data, where in addition to the catch blocks, all normal human body blocks were also regressed out using FIR modeling with 25 predictors per block. A new set of IC was reconstructed from this omitted dataset and the spatial map differences between the original and condition-omitted networks presumably reflect the effect of leaving out human body modulations. Since the estimation of group ICs involves randomization procedures, condition-omitted networks were directly reconstructed from the original aggregated group ICs with GIG-ICA on the new dataset in order to avoid confounds (Du and Fan, 2013).

For the body-selective networks defined above, the difference between the original and conditionomitted maps was computed for each participant and each run. The difference maps were then averaged across runs and entered a group-level t-test against zero and underwent the same cluster-threshold correction. For those human-body-modulated nodes, we expected that their network connectivity would decrease after removing the human body blocks, resulting in lower IC weights in the condition-omitted maps.

## ACKNOWLEDGMENTS

This work was supported by the European Research Council (ERC) FP7-IDEAS-ERC (Grant agreement number 295673; Emobodies), by the ERC Synergy grant (Grant agreement 856495; Relevance), by the Future and Emerging Technologies (FET) Proactive Program H2020-EU.1.2.2 (Grant agreement 824160; EnTimeMent) and by the Industrial Leadership Program H2020-EU.1.2.2 (Grant agreement 825079; MindSpaces).

## SUPPLEMENTARY INFORMATION

**Table S1.**
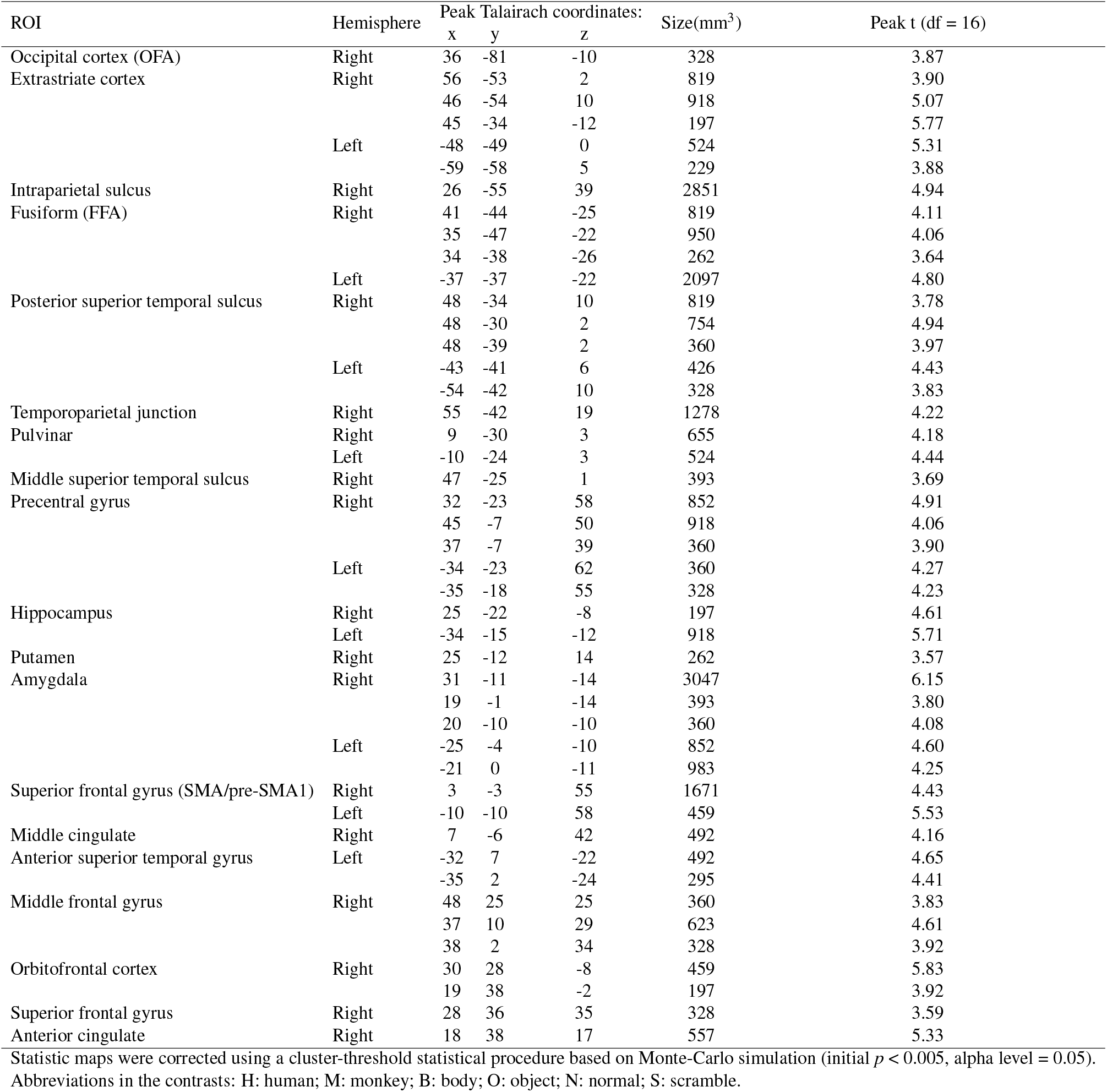
Clusters found by random-effect group GLM. Contrast: human face (normal - scramble) > human object(normal - scramble)

**Table S2.** Clusters found by random-effect group GLM. Contrast: monkey body(normal - scramble) > monkey object(normal - scramble)

| ROI | Hemisphere | Peak Talairach coordinates: |  |  | Size(mm<sup>3</sup>) | Peak t (df = 16) |
| --- | --- | --- | --- | --- | --- | --- |
|  |  | x | y | z |  |  |
| Cuneus | Right | 16 | -76 | 22 | 393 | 4.40 |
|  |  | 13 | -81 | 26 | 360 | 4.41 |
| Extrastriate cortex (EBA) | Right | 46 | -70 | 2 | 6390 | 8.06 |
|  |  | 38 | -60 | 10 | 4653 | 5.28 |
|  | Left | -37 | -62 | 11 | 1180 | 4.61 |
|  |  | -48 | -66 | 12 | 1049 | 4.11 |
|  |  | -44 | -74 | 2 | 229 | 3.84 |
|  |  | -45 | -74 | 12 | 295 | 3.69 |
| Superior parietal lobule | Right | 16 | -70 | 60 | 229 | 4.52 |
|  | Left | -26 | -61 | 60 | 2589 | 5.26 |
| Posterior superior temporal sulcus | Right | 48 | -39 | 10 | 1343 | 4.62 |
| Fusiform (FBA) | Right | 38 | -23 | -14 | 295 | 4.34 |
| Middle superior temporal sulcus | Left | -52 | -13 | -2 | 229 | 3.99 |
| Inferior precentral sulcus (PMv) | Right | 59 | 7 | 17 | 229 | 4.41 |
| Anterior superior temporal gyrus | Left | -46 | 7 | -12 | 295 | 4.95 |
| Superior frontal gyrus | Left | -16 | 38 | 29 | 295 | 5.07 |

Statistic maps were corrected using a cluster-threshold statistical procedure based on Monte-Carlo simulation (initial $p < 0.005$ , alpha level = 0.05).
Abbreviations in the contrasts: H: human; M: monkey; B: body; O: object; N: normal; S: scramble.

